# Effective interferon (IFN)-λ treatment regimen to control lethal MERS-CoV infection in mice

**DOI:** 10.1101/2021.05.26.445685

**Authors:** Ronald Dijkman, Muneeswaran Selvaraj, Hans Hendrik Gad, Rune Hartmann, Sunil More, Stanley Perlman, Volker Thiel, Rudragouda Channappanavar

## Abstract

Effective broad-spectrum antivirals are critical to prevent and control emerging human coronavirus (hCoV) infections. Despite considerable progress made towards identifying and evaluating several synthetic broad-spectrum antivirals against hCoV infections, a narrow therapeutic window has limited their success. Enhancing the endogenous interferon (IFN) and interferon-stimulated gene (ISG) response is another antiviral strategy known for decades. However, the side effects of pegylated type-I IFNs (IFN-Is) and the pro-inflammatory response detected after delayed IFN-I therapy have discouraged their clinical use. In contrast to IFN-Is, IFN-λ, a dominant IFN at the epithelial surface, is shown to be less pro-inflammatory. Consequently, we evaluated the prophylactic and therapeutic efficacy of IFN-λ in hCoV infected airway epithelial cells and mice. Human primary airway epithelial cells treated with a single dose of IFN-I (IFN-α) and IFN-λ showed similar ISG expression, whereas cells treated with two doses of IFN-λ expressed elevated levels of ISG compared to IFN-a treated cells. Similarly, mice treated with two dose IFN-λ were better protected compared to mice receiving a single dose, and a combination of prophylactic and delayed therapeutic regimens completely protected mice from lethal MERS-CoV-infection. A two dose IFN-λ regimen significantly reduced lung viral RNA and inflammatory cytokine levels with marked improvement in lung inflammation. Collectively, we identify an ideal regimen for IFN-λ use and demonstrate the protective efficacy of IFN-λ in MERS-CoV infected mice.

## Introduction

Human CoV infections can cause mild to moderate or severe respiratory illness in humans. Among the hCoVs, seasonal/common-cold hCoV (NL63, 229E, OC43, and HKU1) infections are usually mild and account for approximately 15% of all common cold infections and 10-20% of all respiratory tract infection hospitalization in children (1–3). Emerging high pathogenic CoVs on the contrary cause severe respiratory illness as demonstrated by the high case fatality rates of 35%, 10%, and 2.5% upon MERS-CoV, SARS-CoV, and SARS-CoV-2 infection, respectively (4–11). While SARS-CoV and MERS-CoV each caused approximately 800 deaths, the recently emerged SARS-CoV-2 has infected 144 million people with more than 3.3 million deaths (4, 5, 10, 12). Since humans are immunologically naïve to emerging hCoVs, these viruses rapidly replicate and evade the immune response and cause severe disease, particularly in the elderly and in individuals with co-morbid conditions. Identification of SARS-, MERS-, and SARS-CoV-2-like CoVs in bats and other intermediate hosts highlights the likely spill-over of these viruses from an animal reservoir into the human population, causing future outbreaks (13–15).

Broad-spectrum effective antivirals are urgently needed to control ongoing COVID-19 and any future hCoV or other emerging virus infections. Following SARS-CoV-2 emergence, significant progress has been made towards identifying, developing, and evaluating the therapeutic efficacy of several broad-spectrum and CoV-specific antivirals. The broad-spectrum anti-CoV drugs that completed clinical trials or those in advanced stage of clinical trials include remdesivir (nucleoside analog), EIDD-2810 (nucleoside analog), lopinavir/ritonavir (protease inhibitor), and camostat mesylate (TMPRSS2 inhibitor) (16, 17) (NCT04392219 and NCT0445739) although none of this has demonstrated substantial clinical efficacy. CoV-specific antivirals such as CoV 3CLPro/PLpro protease inhibitors and monoclonal antibodies against spike protein (S) or receptor binding domain of the S protein, are also being investigated for their preclinical and clinical antiviral efficacy (18–20). Several of these broad-spectrum and CoV-specific antivirals effectively reduced viral load in *in vitro* studies (18, 21–24). Preclinical studies in animal models and recent human clinical trials also showed the therapeutic efficacy of these antiviral agents when used in the early stages of infection (21, 24–27). However, so far, antivirals used at later times of infection provided no significant therapeutic benefit over placebo controls, particularly in individuals with severe/critical disease (28–34).

IFNs produced following virus infections facilitate endogenous antiviral state by promoting ISG induction. ISGs reduce virus burden by inhibiting virus entry, replication, and or release (35, 36). Among the three class of IFNs (IFN-α/β, IFN-g, and IFN-λ), IFN-λ is the predominant IFN produced by mucosal epithelium and plays a non-redundant role in host protection during viral lung infections (37–39). Although pegylated (peg-) IFN-α2/ IFN-β is in clinical use for treating chronic virus infections and autoimmune disorders, side effects caused by its chronic use have limited their therapeutic use (40, 41). Our preclinical mouse hCoV studies and recent clinical data from COVID-19 patients also demonstrated the detrimental effect of delayed peg-/recombinant IFN-α/β therapy (42, 43). IFN-λ on the contrary is less pro-inflammatory, and peg-IFN-λ has relatively fewer side effects compared peg-IFN-α/β (44, 45). Consequently, peg-IFN-λ is in clinical trials to treat chronic virus infection and, more recently COVID19 (44–46). Despite recent advances, the prophylactic use of IFN-λ, ideal route administration, and an optimal regimen of IFN-λ to prevent and treat hCoV infections are not well established.

Here, we report that a two dose IFN-λ regimen induces robust ISG expression compared to IFN-α/β as opposed to single-dose IFN-α/βand IFN-λ regimens in primary human airway epithelial cells. We also show that prophylactic and early therapeutic intranasal IFN-λ administration protects mice from lethal hCoV infection, whereas as delayed (relative to virus replication) IFN-λ therapy is detrimental. Of note, a combination of prophylactic and delayed therapeutic regimen provided complete protection from lethal hCoV infection. Our results collectively demonstrate an ideal treatment regimen and the protective role of IFN-λ upon lethal hCoV challenge in mice.

## Material and Methods

### Human airway epithelial cells

Primary human airway epithelial cells were procured from patients who underwent surgical lung resection in their diagnostic pathway for any pulmonary disease and who gave informed consent. This was done in accordance with our ethical approval (EKSG 11/044, EKSG 11/103 and KEK-BE 302/2015). Pathologically examined tracheobronchial segments were used as starting material for isolation, propagation, and establishment of well-differentiated primary human airway cell (hAEC) cultures (47). All hAEC cultures were allowed to differentiate for >3 months.

### Viruses and recombinant Interferons

Human coronavirus 229E, strain Inf-1, was generated and rescued using the vaccina virus reverse genetic system as described previously, and propagated and titrated on Huh7 cells (47). The Middle East respiratory syndrome virus, strain EMC, was isolated from patient material and further propagated and titrated on Vero-B4 cells (48). The mouse-adapted MERS-CoV was generated and titrated as described elsewhere (49). The recombinant human interferon lambda 3 (IFN-λ3) and mouse IFN-λ2 were produced as described previously (38, 50). The mammalian universal Type I interferon (IFN-αA/D) was procured commercially (Sigma-Aldrich; I4401) and recombinant murine IFN-λ2 was purchased from Peprotech (Catalog # 250-33).

### IFN treatment and Trans Epithelial Electrical Resistance (TEER) measurement

Well-differentiated hAEC cultures were stimulated for 24 hours with either IFN-λ3 (100ng/ml) or IFNa A/D (100U/ml) or were left untreated. Medium was exchanged every 24 hours and were indicated supplemented with either IFN-λ3 (100ng/ml) or IFNa A/D (100U/ml). The barrier integrity was qualitative assessed through measurement of the Trans Epithelial Electric Resistance (TEER) using an EVOM2 instrument (World Precision Instrument). Prior to the experimental endpoint, 100uL of 0.9% sodium chloride saline solution, supplemented with 1.25 mM CaCl2 and 10 mM HEPES (TEER solution), was applied to the apical surface of each hAEC cultures. Cell cultures were allowed to equilibrate for 10 minutes at 37°C, followed by TEER measurement using the electrode chopsticks. In between measurements, the chopstick electrode was washed and sterilized through subsequent washing steps in TEER solution and 70% EtOH. Control cells were mock treated.

### Mice infections and morbidity studies

12-14-week male human dipeptidyl peptidase-knock-in (hDPP4-KI) mice were treated with one or two doses of recombinant mouse IFN-λ-2 (1.5 to 2.0μg/mouse, intranasal in PBS) at the indicated time points. Mice were challenged with 1000 PFU of mouse-adapted MERS-CoV (49), and were monitored for morbidity and mortality. Mice were weighed and monitored daily for clinical illness. Mice weight loss of >30% of the initial body weight was considered as endpoint and mice were humanly euthanized as per American Veterinary Medical Association (AVMA) guidelines.

### Gene expression analysis of IFN-treated hAEC cultures and MERS-CoV infected lungs

Total cellular RNA from IFN-treated hAEC cultures was extracted with the NucleoMag RNA kit (Macherey-Nagel) according to the manufacturer’s guidelines on a Kingfisher Flex Purification system (Thermofisher). Reverse transcription was performed with Moloney murine leukemia virus reverse transcriptase (MMLV-RT) according to the manufacturer’s protocol (Promega) using 200 ng of total RNA. Two microliters of tenfold diluted cDNA were amplified using Fast SYBR^™^ Green Master Mix (Thermofisher) according to the manufacturer’s protocol using primers targeting 18S and MxA as described previously (48). Measurements and analysis were performed using an ABI7500 instrument and software package (Thermofisher). Relative gene expression was calculated using the 2^-ΔΔCt^ method (51) and is shown as fold induction of compared to that of untreated controls. For quantification of viral RNA, apical washes of Human coronavirus 229E (HCoV-229E) and MERS-CoV-infected hAECs were extracted using the NucleoMag VET (Macherey-Nagel) according to manufacturer’s guidelines on a Kingfisher Flex Purification system (Thermofisher). Measurements and analysis were performed as described previously (48).

Control and hCoV infected mouse lungs were collected in Trizol at different days post-infection. Isolated RNA (TriReagent, Ambion) was DNAse digested to eliminate genomic DNA contamination (RQ1 DNAse kit, Promega) as per manufacturer’s instructions. cDNA synthesized with MMLV-RT (Life Technologies) was used to quantitate genomic viral RNA and mRNA levels of cytokines and chemokines (Power-Up SYBR-green master mix, Life Technologies) in a Quant-Studio 5- or 6-flex.

### Histology Studies

Animals were anesthetized and transcardially perfused with 10 mL PBS followed by 5 mL zinc formalin. Lungs were removed, fixed in zinc formalin, and paraffin embedded. Tissue sections were stained with hematoxylin and eosin (H & E) and examined and scored by light microscopy in a blinded fashion by board certified veterinary pathologist as described previously (52). Control and rIFN-λ–treated lungs were scored for edema with scores of 0, 1, 2, 3, and 4 representing lung areas with 0%, less than 3%, 6%–33%, 33%–66%, and more than 66% detectable proliferation, respectively. Lungs were also scored for inflammatory cell infiltration, with scores of 0, 1, 2, 3, and 4 representing areas with 0%, less than 3%, 6%–33%, 33%–66%, and more than 66% of perivascular poly/mononuclear cell distribution, respectively.

### Statistical analysis

Results were analysed using Student’s t test. Data in bar graphs or bar/scatter plots are represented as mean +/-SEM. Weight loss and survival curves were assessed for statistical significance using log-rank (Montel-Cox test) or Gehan-Breslow-Wilcoxon test. *P<0.05, **P<0.01, or ***P<0.001 was used to determine statistical significance.

## Results

### Two dose IFN-λ regimen induces robust ISG expression in human airway epithelial cells

Epithelial cells express both IFN-α/β receptor (IFNAR) and IFNλR (39, 53–55). Therefore, we first assessed whether airway epithelial cells induce differential levels of ISGs when stimulated with similar amounts of recombinant human (rh)IFN-α/βand rhIFN-λ (48, 56). Isolated primary hAECs were treated with single or two-dose regimen of 100U/ml of IFN-α/β or 100ng/ml of IFN-λ. At different days post-treatment, airway epithelial cells were collected to assess mRNA levels of ISGs (i.e., MxA). As shown in figure 1A, MxA expression was similar in single dose IFN-λ and IFN-α/βtreated groups at different days post-treatment. Interestingly, MxA expression was significantly higher in cells that received two doses of rhIFN-λ compared to two dose rhIFN-α treated cells. IFNs alter the epithelial cell barrier during virus infections. To assess if single or dual treatment with IFNa or IFN-λ affects barrier integrity, we measured trans epithelial electrical resistance (TEER) at different days post-IFN-treatment. Our result showed no significant difference in TEER in control and rhIFN treated or between rhIFN-a and rhIFN-λ treated cells at different days post-treatment. These results demonstrate that the dual IFN-λ treatment regimen induces a significantly elevated level of ISG expression in airway epithelial cells without affecting the barrier integrity.

**Figure 1:**
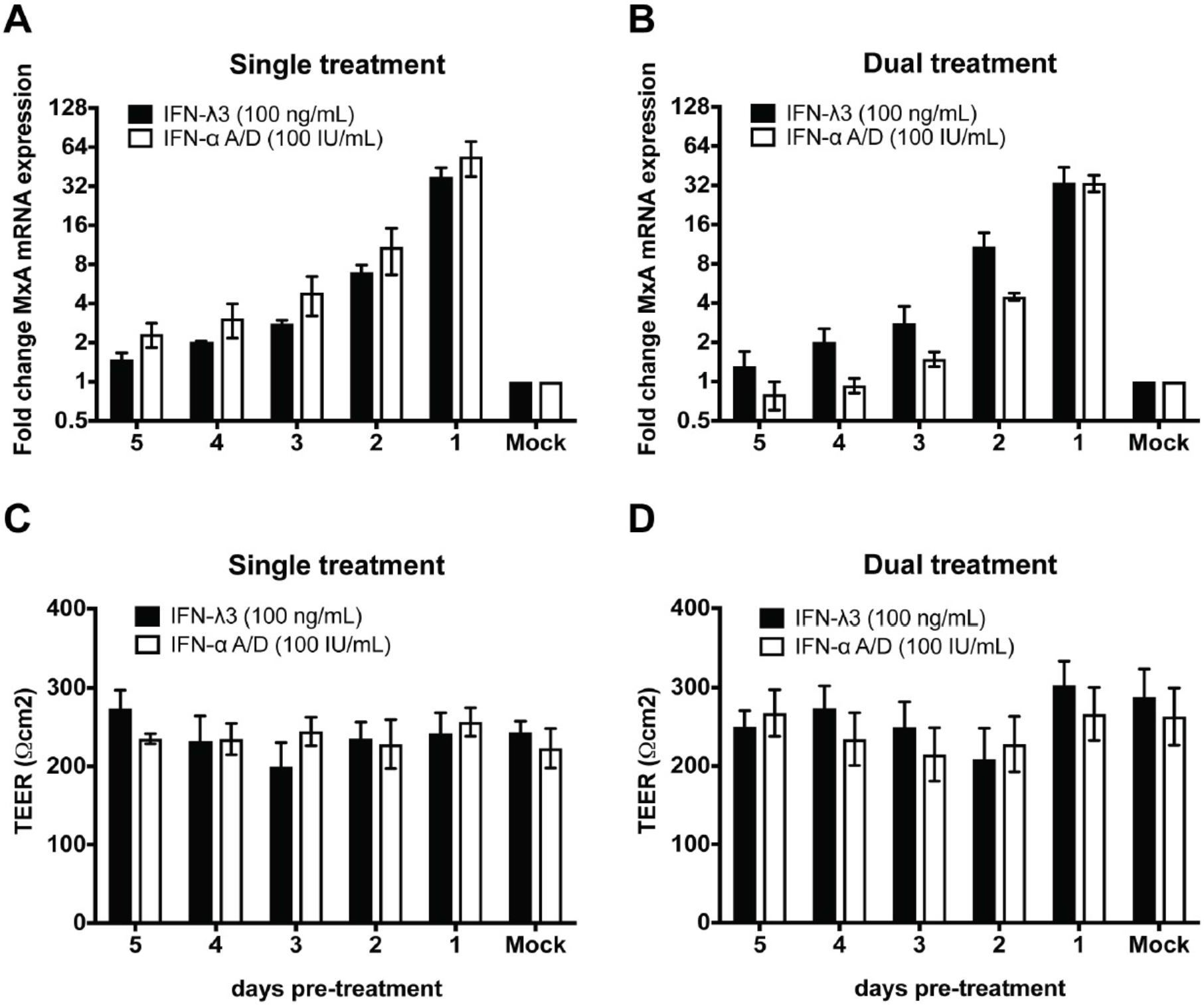
Two dose IFN-λ regime induces robust ISG expression in hAECs. To determine the duration of the antiviral response, well-differentiated hAEC cultures were treated with a single or double dose of IFN-aA/D (100IU/ml) or hIFN-λ 3 (100 ng/ml) at indicated time points prior to expression analysis of MxA mRNA. For the two-dose regime hAEC cultures were stimulated 6 days prior to readout, followed by a restimulation at the indicated time points. The fold change in MxA mRNA expression over mock is shown for the individual **A)** single or **B)** two dose IFN-λ regimes. In addition to the duration of the antiviral response, the epithelial barrier integrity among the **C)** single and **D)** two dose regimes was assessed by measuring the transepithelial electrical resistance (TEER) of each regime prior to whole cell lysis. Data is shown as mean and standard deviation from 3 biological donors with two technical replicates.

### Two dose IFN-λ treatment reduces hCoV titers in human airway epithelial cells

We next examined whether increased ISG expression in dual IFN-λ treatment groups correlates with reduced virus titers. Well-differentiated hAEC cultures were treated with single or two doses of rIFN-λ (100ng/ml or 100 IU/ml) before and or after infection with hCoV-229E (4000 PFU) or MERS-CoV (4000 PFU) (Figure 2A-D). Cell apical washes collected at 72 hours post-infection were used to assess virus load. Our results show that single-dose IFN-λ treatment reduced HCoV-229E genomic RNA levels compared to mock-treated cells, with treatment on one-day before (−1d) and on the day of infection (0d) being highly effective (Figure 2A) in suppressing viral RNA levels. Compared to single-dose regimen, two-dose IFN-λ treatment further reduced hCoV-229E RNA levels (Figure 2B) demonstrating that two-dose regimen is much more effective than the single-dose treatment. We then evaluated the efficacy of rhIFN-λ in suppressing MERS-CoV replication. As shown in Figure 2C-D, MERS-CoV RNA levels were reduced upon IFN-λ administration compared to mock treatment, with two-dose regimen being more effective than one dose IFN-λ in reducing viral RNA levels (albeit not statistically significant). Nonetheless, we show that IFN-λ treatment reduced hCoV RNA levels compared to controls, and hCoV-229E was highly sensitive to IFN-λ-treatment compared to MERS-CoV in hAEC cultures.

**Figure 2:**
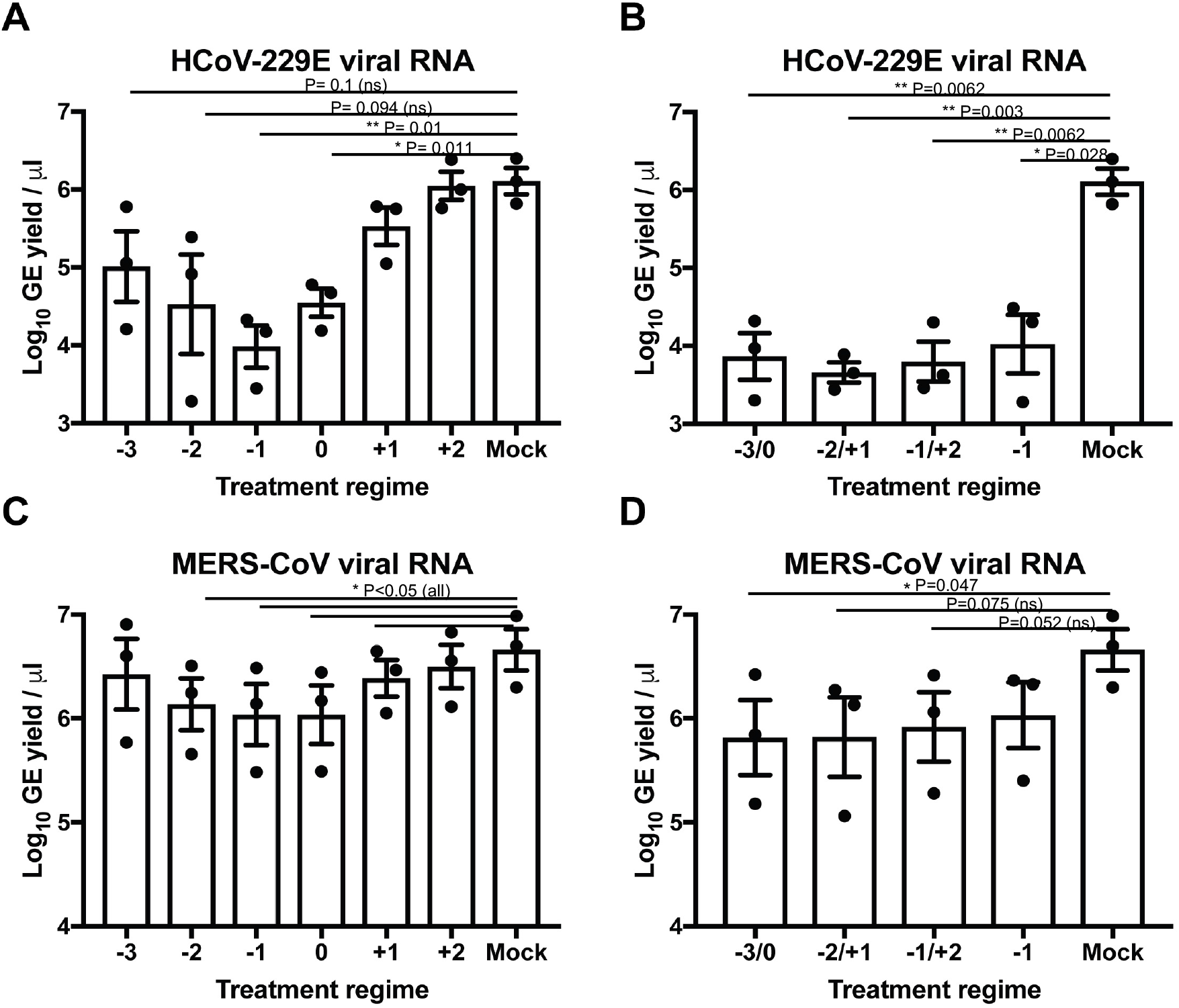
IFN-λ treatment reduces hCoV replication in human airway epithelial cells. Well-differentiated hAEC cultures were treated with a single or two doses of IFN-aA/D (100IU/ml) or hIFN-λ 3 (100 ng/ml) at indicated time points (minus (-) symbol refers to day-before infection, and plus (+) to days-after infection). The hAEC cultures were challenged with 4000 PFU of HCoV-229E or MERS-CoV and 72 hours post-infection the viral RNA load (Genome Equivalents (EQ) / uL) was measured by qPCR. Bar graphs represent HCoV-229E **(A, B)** or MERS-CoV **(C, D)** genomic RNA levels in control and IFN-λ treated hAEC cultures during a single or double dose regime, respectively. Data is shown as mean and standard deviation from 3 biological donors with two technical replicates. Statistical significance was determined using Student’s *t* test with * P<0.05 and ** P<0.01, *** P<0.001, and **** P<0.0001.

### Prophylactic and combination of prophylactic plus therapeutic IFN-λ provides protection

Recent studies from others and our laboratory showed that timing of IFN-α/β therapy is a critical determinant of host protection versus pathology (42, 43, 57). While prophylactic and early IFN-α/β administration protected mice from lethal hCoV disease, delayed/late (relative to virus replication) IFN-α/βtherapy caused severe disease. Since IFN-λ is less pro-inflammatory, we hypothesized that the therapeutic administration of IFN-λ would provide better protection compared to IFN-α/β. To test this possibility, hDPP4-KI mice were treated with rmIFN-λ before or after lethal MERS-CoV infection. Our results showed that single IFN-λ instillation before (−1 and −2 dpi), on the day (0 dpi), and early (+1dpi) after MERS-CoV-MA infection protected mice from death but afforded only partial protection from weight loss as compared to PBS-treated controls (Figure 3A and B). Conversely, delayed IFN-λ treatment (+2dpi) was detrimental (Figure 3A) in comparison to early IFN-λ and PBS treated mice. We then examined whether a combination of pre- and post-infection (i.e., delayed) IFN-Λ treatment regimen would further protect mice from morbidity and weight loss. hDPP4-KI mice were treated with IFN-λ at various days before and or after infection with a lethal dose of MERS-CoV-MA. As shown in Figure 3B, our results showed complete protection in hDPP4-KI mice treated with IFN-λ before and after lethal MERS-CoV-MA infection, including mice that received a second dose of IFN-λ at +2dpi. Together, our results demonstrate that a) prophylactic IFN-λ therapy is protective during MERS-CoV infection, and b) while a single dose of delayed IFN therapy is pathogenic, an IFN-λ combination regimen that includes treatment before and after infection protects the host from lethal MERS-CoV challenge.

**Figure 3:**
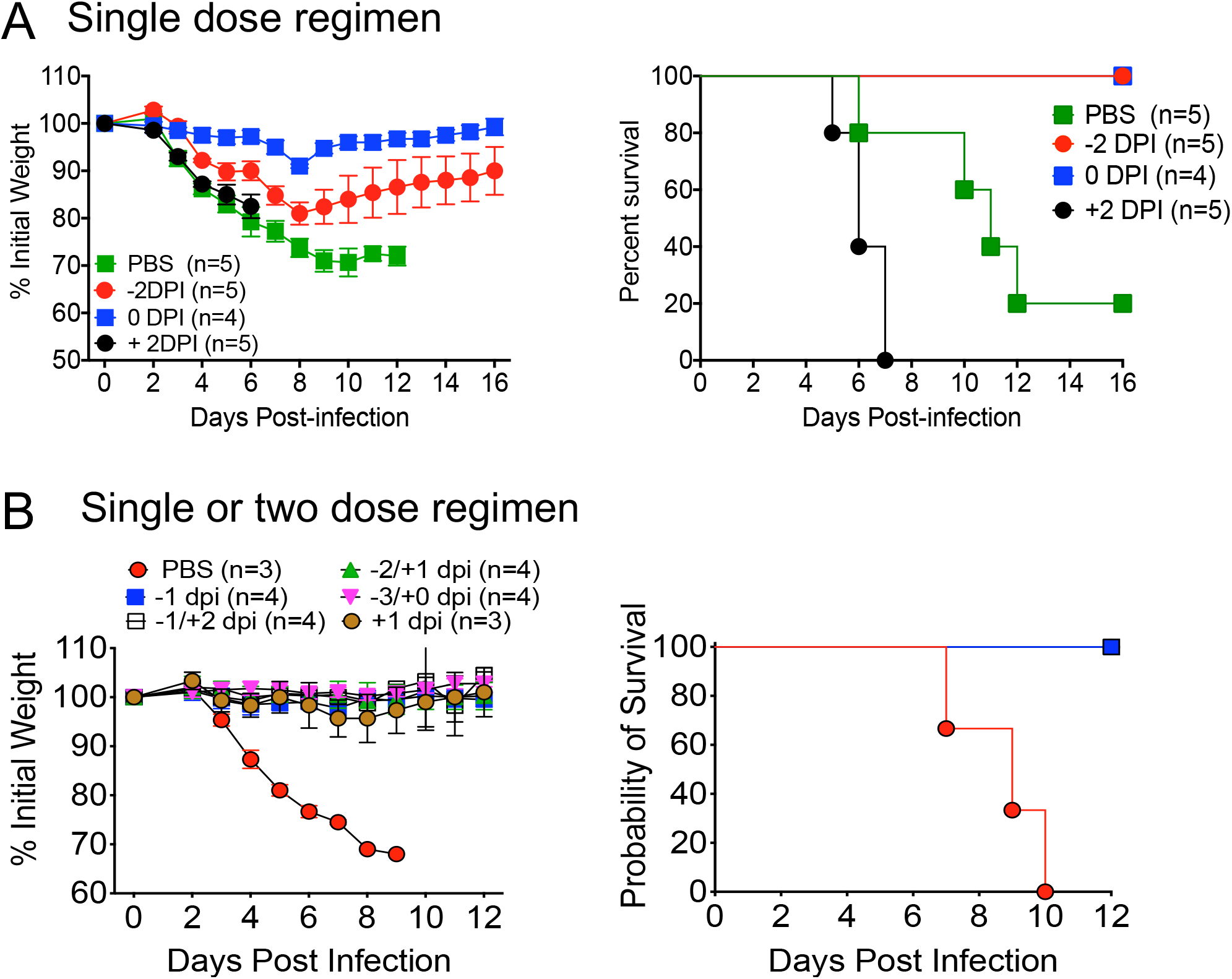
Optimal prophylactic and therapeutic regimen of IFN-λ provides protection from lethal hCoV infection. Male hDPP4-KI mice were treated with a single or two doses of mIFN-λ (1.5 to 2.0μg/mouse, vial intranasal route) at indicated time points (minus (−) symbol refers to day-before infection, and plus (+) to days-after infection). Mice were challenged with 1000 PFU of mouse-adapted MERS-CoV and mice were monitored for morbidity and mortality for 14 days. **A)** Shows weight loss and survival in mice receiving single dose of rmIFN-λ at indicated days pre- or post-infection. **B)** Weight loss and survival curves show morbidity and mortality in mice receiving single or two regimens of rmIFN-λ. The data are derived from one (A) or are representative of two experiments (B) with 3-5 mice in each group per experiment.

### Combination IFN-λ regimen reduces viral RNA and inflammatory cytokine levels upon MERS-CoV infection

IFN response timing determines inflammatory response upon hCoV infection. An early IFN response/therapy leads to controlled inflammation, whereas a delayed/late response facilitates exaggerated inflammation (42, 43). To test whether delayed single and dual IFN-λ therapy has differential effects on virus replication and ISG/cytokine production, we evaluated MERS-CoV-RNA, ISG, and inflammatory cytokine levels in PBS-, delayed-single-, and prophylactic plus therapeutic IFN-λ treated MERS-CoV infected lungs. As shown in Figure 4A, dual IFN-λ treatment regimen significantly reduced MERS-CoV-RNA levels compared to PBS treated and delayed single dose IFN-λ treated mice when evaluated at 4 dpi. Notably, we did not observe any difference in MERS-CoV-RNA levels between PBS and delayed IFN-λ treated mice (Figure 4A). Further examination of MERS-CoV infected lungs showed reduction in ISG (CXCL-10 and ISG15) and inflammatory cytokine levels (CXCL-1, IL-6, TNF, and IL-1b) in the lungs of prophylactic plus therapeutic IFN-λ treated mice in comparison to PBS and delayed single dose IFN-λ treatment. In contrast, no significant difference in ISG or inflammatory cytokine levels was observed between PBS and delayed single-dose IFN-λ treated mice. These results collectively show that the two-dose IFN-λ regimen effectively reduces MERS-CoV replication (as measured by viral RNA levels) and ISG and inflammatory cytokine expression, likely protecting the host from lethal MERS.

**Figure 4:**
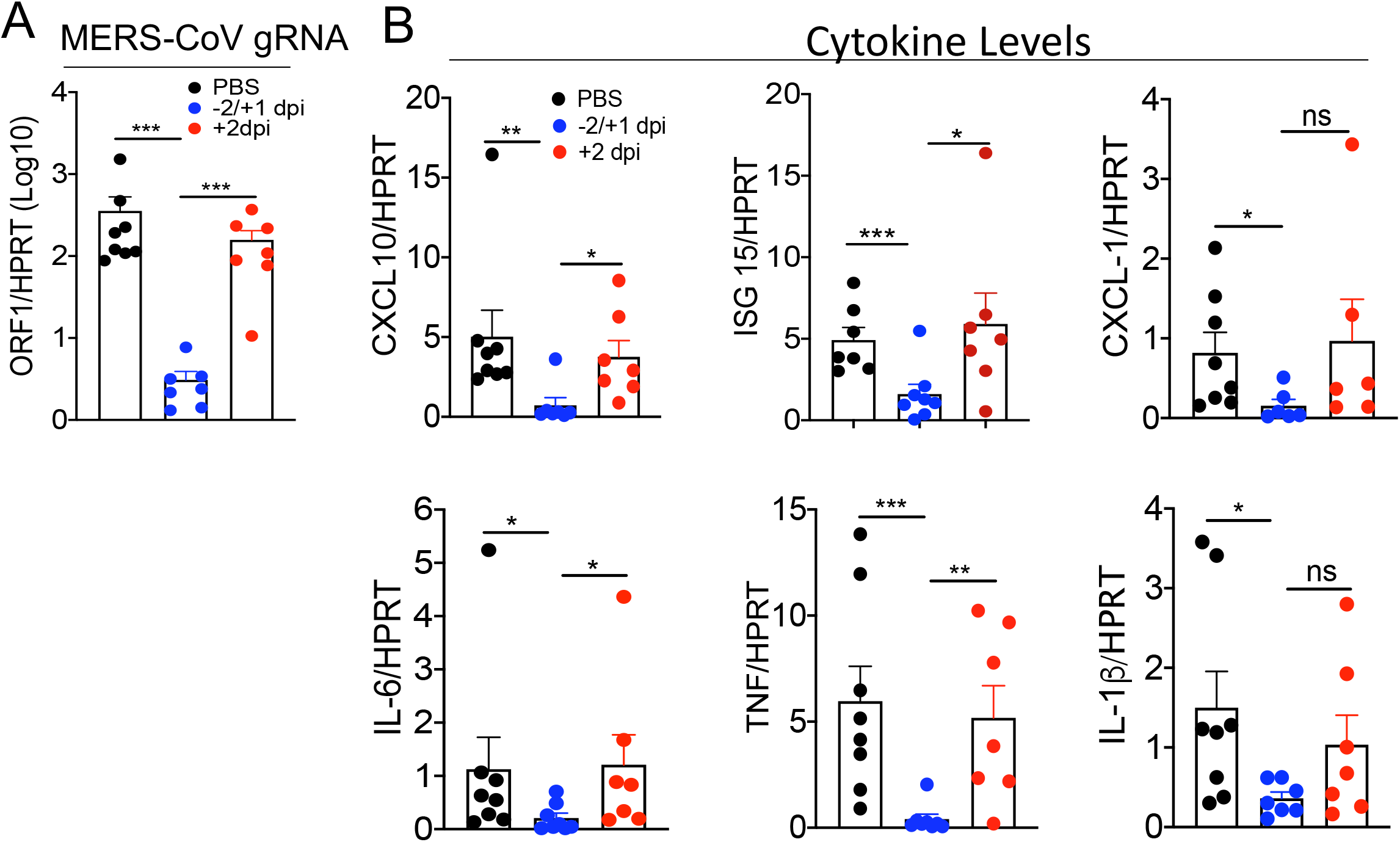
Prophylactic plus therapeutic IFN-λ regimen effectively reduces hCoV titers. Male hDPP4-KI mice were treated with a single or two doses of mIFN-λ (1.5 to 2.0μg/mouse, vial intranasal route) at indicated time points (minus (-) symbol refers to day-before infection and plus (+) to days-after infection). Mice were challenged with 1000 PFU of mouse-adapted MERS-CoV and mice were monitored for morbidity and mortality for 14 days. **A)** Bar graphs represent MERS-CoV genomic RNA levels in control and IFN-λ treated lungs at 4 DPI, with each scatter points representing a mouse. **B)** Bar graphs represent mRNA levels of ISGs and inflammatory cytokines in control and IFN-λ treated lungs at 4 DPI, with each scatter points representing a mouse. Data are pooled from 2 experiments with 3-4 mice per group per experiment. Statistical significance was determined using Student’s *t* test with * P<0.05 and ** P<0.01.

### IFN-λ treatment provides protection from MERS-CoV induced lung pathology

To assess the lung pathology in control PBS treated and IFN-λ treated mice, we collected PBS-perfused lungs at 4dpi from control-PBS and IFN-λ treated mice infected with a lethal dose of MERS-CoV. Histological examination of control PBS- and delayed (+2dpi) IFN-λ-treated lungs showed significantly increased alveolar and bronchial edema and fibrin deposition, associated with mild to moderate accumulation of mononuclear cells and neutrophils in the perivascular spaces, alveoli, and interstitium (Figure 5A-B). In contrast, lungs from mice receiving prophylactic plus therapeutic (−2 and +1 dpi) and early therapeutic IFN-λ (+1dpi) showed significant reduction in the pulmonary edema with moderately reduced inflammatory cell infiltration as compared to PBS controls (Figure 5A-B).

**Figure 5:**
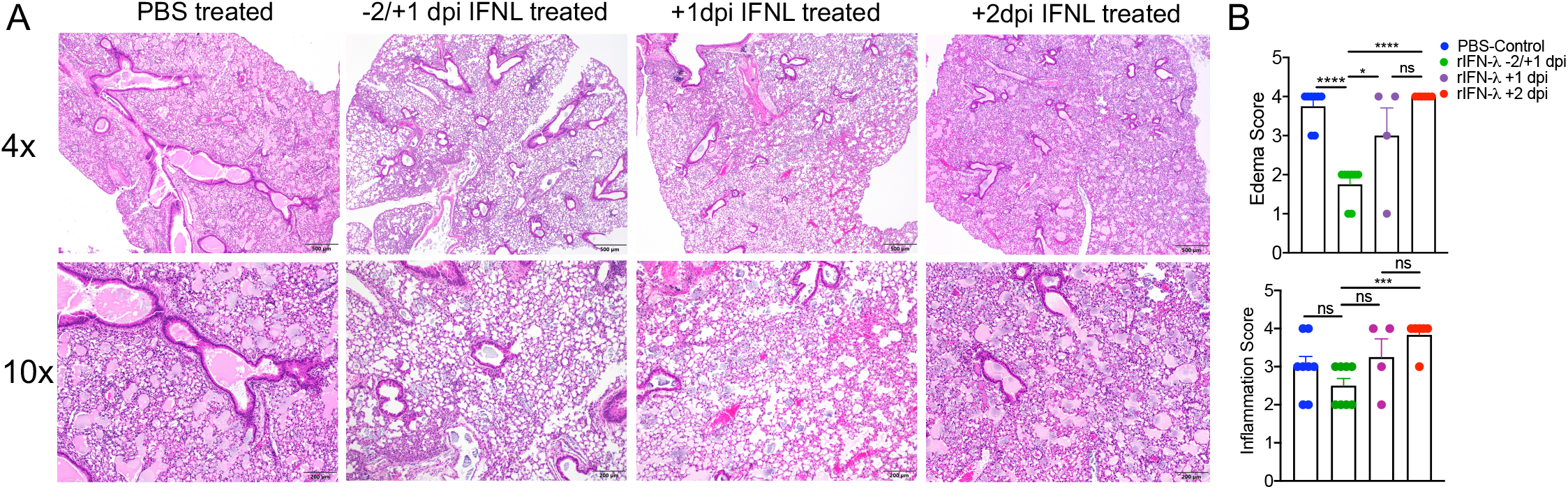
IFN-treatment reduces MERS-CoV-induced lung pathology. Control PBS and rIFN-λ treated (at different days before and or after infection) mice were euthanized at 4 dpi. PBS-perfused lungs collected in Zn-formalin were H & E stained for assessing lung pathology. A) Histology pictures represent lung edema, inflammation, and epithelial cell damage. (B) Scatter plot bar graphs show average edema (top panel) and inflammation (bottom panel) in control and rIFN-λ treated lungs. Each scatter plot represents individual animal. Histology pictures are representative 1-2 experiments with 3-4 mice per group per experiment. Statistical significance was determined using Student’s *t* test with *** P<0.0003 and **** P<0.0001.

## Discussion

The role of IFN-λ during hCoV infection and its therapeutic potential is beginning to be elucidated. Recent COVID19 studies show IFN deficiency in patients with severe disease, while others demonstrate an association of elevated IFN signatures in patients with critical COVID19 illness (58–62). Results from recent COVID19 clinical trials demonstrate the therapeutic advantages of IFN-λ in reducing the SARS-CoV-2 burden in COVID19 outpatients (46). However, the protective efficacy of prophylactic and early- and delayed-therapeutic potential of IFN-λ is not well understood. Consequently, we evaluated the prophylactic and therapeutic efficacy and an optimal treatment regimen of IFN-λ in mice infected with a lethal dose of hCoVs. We show significant protection from lethal MERS-CoV infection in young mice following prophylactic IFN-λ administration provides. We also demonstrate that while an early therapeutic IFN-λ instillation is protective, a delayed treatment is detrimental. Of note, a combination of prophylactic and delayed therapeutic administration of IFN-λ protected mice from severe hCoV induced disease.

Type I and type III IFNs play a critical role in host defence against a variety of virus infections (35, 37, 63). IFN-I is produced by the majority of virus-infected cells, with dendritic cell and monocyte-macrophage subsets specialized in making large quantities of IFN-α/β. IFNAR is ubiquitously expressed, as a result IFN-α/β induce an antiviral state in all nucleated cells (63). IFN-λ on the contrary, is produced by epithelial cells and a subset of dendritic cells (37, 39). Given that IFNλR expression is limited to mucosal epithelial cells, DCs, and neutrophils, the IFN-λ mediated antiviral responses are limited to these cells. IFN-λ/IFNλR signalling induces robust ISG but reduced inflammatory response in comparison to IFNAR signaling (64–66). Consequently, IFN-λ is recommended for therapeutic use during acute and chronic virus infections. Recent studies demonstrate that IFN-λ is the dominant interferon that plays a non-redundant role in host protection during influenza A virus (IAV) infection (37, 39). Upon influenza virus and several enteric virus infections, IFN-λ is expressed at significantly high levels than IFN-α/β (37, 39, 67). Similarly, hCoVs infected airway epithelial cells express increased levels IFN-λ compared to IFN-α/β (68). Mice infected with hCoVs such as SARS-CoV, MERS-CoV, and SARS-CoV-2 also express similar or elevated level of IFN-λ in the lungs (43, 69). Conversely, intranasal instillation of hepato/neurotropic mouse coronavirus (MHV-A59) showed reduced lung IFN-λ levels compared to IFN-α/β (Qing H *et al*., 2020). Overall, these results suggest that IFN-λ is likely the dominant IFN produced at the respiratory mucosa during hCoV infections.

We recently showed that the protective efficacy of endogenous and exogenous IFN-α/β is dependent on the timing of IFNs response relative to virus replication (42, 43). Our results further showed that the detrimental role of delayed IFN response is mediated by the excessive inflammatory response, in part caused by monocyte-macrophage and neutrophil-mediated exaggerated inflammatory cytokine/ chemokine responses (43). IFN-λ on the contrary is shown to be less pro-inflammatory due to limited expression of IFN-λR on inflammatory cells (65, 66) and, as a result, is believed to provide better protection as opposed to IFN-α/β following therapeutic administration. Our results, however, showed that delayed IFN-λ administration is pathogenic (Figures 3 and 6), which is similar to outcomes resulting from delayed IFN-α/β treatment. As shown in Fig 3, a combination of prophylactic and delayed therapeutic antiviral therapy will overcome these detrimental effects. However, given that IFN-λ is less pro-inflammatory, whether disease exacerbation following delayed IFN-λ administration occurs due to robust inflammation (43) or epithelial cell apoptosis (70, 71) resulting in impaired tissue repair requires further investigation. Similar to results from the timing of IFN-α/β response, we observed efficient virus clearance upon early but not delayed IFN-λ treatment (Figure 5). Such an impaired virus clearance is also observed upon delayed administration of other anti-CoV antivirals such as remdesivir, and anti-S and anti-S-RBD monoclonal antibodies (17, 33, 34, 72–74), suggesting that early antiviral therapy is critical to effectively reduce virus burden and protect the host from lethal disease.

In addition to an optimal IFN-λ regimen to protect the host from lethal virus infection, route of administration likely plays a key role in the disease outcome. IFNs are mainly administer via the subcutaneous (s/c) or intravenous route to treat virus infections and other immunoinflammatory conditions. Considering hCoV replication is largely limited to airways and lungs, and IFN-λR is mainly expressed on epithelial cells, IFN-λ intranasal spray will likely be more effective than the s/c or systemic administration. Furthermore, considering the combination of prophylactic and therapeutic regimen protects mice from lethal MERS-CoV infection, such a treatment regimen would be beneficial in protecting susceptible immunocompromised, aged, and individuals with co-morbid conditions from lethal disease caused by respiratory virus infections. Collectively, our results demonstrate that prophylactic and prophylactic plus therapeutic administration of IFN-λ protects mice from lethal hCoV infection and therefore has the potential for further clinical testing for use in humans to provide protection from emerging and re-emerging viruses.

## Author contribution

RC, RD, SP, and VT conceptualized the study; RC, RD, MS carried out experiments; HHG and RH provided reagents; SM carried out histology evaluation; RC and RD wrote manuscript, and RC, SP, RD, MS, and VT reviewed and edited the manuscript.

## Acknowledgements

This work is supported in part by institutional research fund to RC from Oklahoma State University-Stillwater. Part of the work was conducted at the University of Tennessee Health Science Center-Memphis. Part of this work is also supported by NIH P01 AI060699 (SP) and RO1 AI129269 (SP).

